# Maternal Obesity Decreases Offspring Lifespan

**DOI:** 10.64898/2026.02.04.703858

**Authors:** Eric Moore, Selvarani Ramasamy, Kavitha Kurup, Michael Chan, Mani Saminathan, Natesan Pazhanivel, Kai Ding, Alexandra Ford, Brianne M. Taylor, Karen Jonscher, Arlan Richardson, Jacob E. Friedman, Archana Unnikrishnan

## Abstract

Data in mice, nonhuman primates, and in humans demonstrate that exposure to maternal obesity increases the risk of multiple diseases in offspring. However, little is known about the aging effects of maternal obesity on the offspring. This study shows that maternal obesity significantly reduced the lifespan of both male and female mice born to obese dams despite being weaned onto a healthy diet at three weeks of age. This reduction in longevity was linked to an increase in age-related fibrotic pathology across multiple organs, e.g., liver, heart, and kidney. Gompertz analysis of the lifespan data showed that maternal obesity offspring have reduced lifespan due to detrimental changes established early during development rather than factors that modify aging later-in-life. These findings are translationally significant as they demonstrate that the growing prevalence of MO may lead to a decrease in overall lifespan and increase in age-related diseases in the next generation.

## 1. Introduction

Maternal exposures, including maternal overnutrition, have adverse effects on offspring health and disease risk that persists into adult life, possibly leading to changes in organ structure, function, and metabolism that has the potential to impact aging^1, 2^. The incidence of maternal obesity has dramatically increased in the past decade with nearly half (47%) of pregnancies in North America occurring in women who are overweight/obese^3^. Maternal obesity increases the risk of numerous adverse outcomes on the developing fetus, including premature birth, still birth, and congenital anomalies, as well as health effects that emerge in offspring during childhood and young adulthood (∼25-30 years of age), such as diabetes^4^, liver disease^5^, cardiovascular disease^6, 7^, and cognitive deficits, including behavior problems^4, 8^. However, essentially nothing is known about the long-term effects of maternal obesity later in life. For example, does maternal obesity accelerate aging in offspring and lead to an increase in the occurrence of age-related diseases?

Given the inherent challenges of conducting human aging studies based on maternal obesity, and the difficulty in delineating the lifelong effects of obesogenic diet exposure in both mothers and offspring, animal models become critical. These models are necessary to accurately elucidate the long-term impacts of *in utero* exposure to maternal obesity on offspring as they age *without* the confounding variable of continuous obesogenic diet intake. Studies with rodents and non-human primate models show that offspring born to obese mothers exhibit adverse effects that persist into young adulthood (3 to 6 months of age in rodents), including insulin resistance^9^, hypertension^10^, cardiovascular disease^11^, liver disease^5, 10^, and cognitive decline^12^. Young/adult maternal obesity offspring also show an increase in hallmarks of aging, (e.g., elevated ROS production^13^, DNA damage^14^, inflammation^13, 14^, mTOR signaling^15, 16^, and mitochondrial dysfunction^17^), suggesting that maternal obesity offspring might show accelerated aging. To determine the impact of maternal obesity on aging, we measured the two hallmarks of aging, lifespan and pathology, in mice born to obese dams.

## 2. Results

### 2.1 Offspring Body Composition

Maternal obesity offspring were generated as shown in Fig. S1A from C57BL/6J dams fed either a control diet (standard chow diet) or a high-fat, high-sucrose, western diet for 8 weeks prior to mating. The breeder females fed the western diet showed a significant 20% increase in total body weight and a ∼2.5-fold increase in percent fat mass (%fat) (Fig S1C). At 3 weeks of age, male and female offspring of both groups of dams were weaned to a non-obesogenic control diet which was continued for the rest of their life. Because maternal obesity offspring themselves are more prone to obesity^13^ and because obesity might impact aging^18^, we measured the body composition of the two groups of mice every 6 months until 24-months of age. As shown in Fig. S2, we found no statistically significant difference in either body weight, lean mass, or fat mass in maternal obesity offspring over their lifespan.

### 2.2 Offspring Lifespan

The lifespans of the male and female maternal obesity and control (offspring born to normal weight dams) offspring are shown in Fig. 1A, and the analyses of the lifespan data are presented in Table 1. Maternal obesity significantly reduced the lifespan of both male (p < 0.001) and female (p < 0.0005) mice compared to control mice whether measured by the non-parametric mantel-cox test or the parametric models with Gompertz distribution. Mixed effects Cox proportional hazards analysis also showed that maternal obesity significantly (p < 0.02) increased the hazard rate (i.e., increased instantaneous risk of mortality) in both male and female maternal obesity offspring. A significant reduction of ∼19% in mean (p < 0.01) and median (p < 0.03) lifespan was observed in male maternal obesity offspring. In the female maternal obesity offspring, the mean lifespan was reduced ∼20% (p < 0.0003) but the 8% decrease in median lifespan was not significant. Both male and female maternal obesity offspring showed a ∼10% reduction in the 90^th^ percentile lifespan, which was statistically significant for males (p < 0.006), but not quite significant for females (p = 0.06). Using the maximum lifespan test developed by Gao et al.^19^ to statistically test for significant differences in the upper tail of the distribution in the survival data, we found the maximum lifespan test was significantly reduced for the maternal obesity male offspring (p < 0.006), but again not quite significant for maternal obesity female offspring (p = 0.06).

**Table 1:** Analysis of lifespan data from control and maternal obesity mice.

|  | **Males** |  | **Females** |  |
| --- | --- | --- | --- | --- |
| **Groups** | **control** | **maternal obesity** | **control** | **maternal obesity** |
| **N** | 64 | 50 | 54 | 42 |
| **Mean (± SEM)** | 859 ± 41 | 698 ± 50 (-19%) | 833 ± 23 | 670 ± 39 (-20%) |
| **p Value (Mean)<sup>a</sup>** | 0.01 |  | 0.0003 |  |
| **Median** | 962 | 815 (-15%) | 836 | 765 (-8%) |
| **p Value (Median)** | 0.03 |  | 0.2 |  |
| **90th%** | 1165 | 1051 (-10%) | 1008 | 926 (-8%) |
| **p Value (90th%)<sup>b</sup>** | 0.03 |  | 0.06 |  |
| **Max lifespan test (p Value)<sup>c</sup>** | 0.006 |  | 0.06 |  |
| **Max Lifespan<sup>d</sup>** | 1256 | 1152 (-8%) | 1118 | 1004 (-10%) |
| **Cox Hazard Ratio (95% CI)<sup>e</sup>** | 3.219 (1.249-8.293) |  | 2.751(1.375-5.503) |  |
| **p Value (Cox Hazard Ratio)<sup>e</sup>** | 0.02 |  | 0.004 |  |
| **Mantel-Cox<sup>f</sup> (p-value)** | 0.0002 |  | 0.0005 |  |
| **Gehan-Breslow-Wilcoxon<sup>g</sup> (p-value)** | 0.001 |  | 0.0005 |  |
| **Gompertz a value<sup>h</sup>** | 0.026463 | 0.069183 | 0.008197 | 0.039955 |
| **p Value (Gompertz a)<sup>i</sup>** | <0.0001 |  | <0.0001 |  |
| **Gompertz b value<sup>h</sup>** | 0.003041 | 0.002442 | 0.004685 | 0.003412 |
| **p Value (Gompertz b)<sup>i</sup>** | <0.0001 |  | <0.0001 |  |

**Fig. 1:**
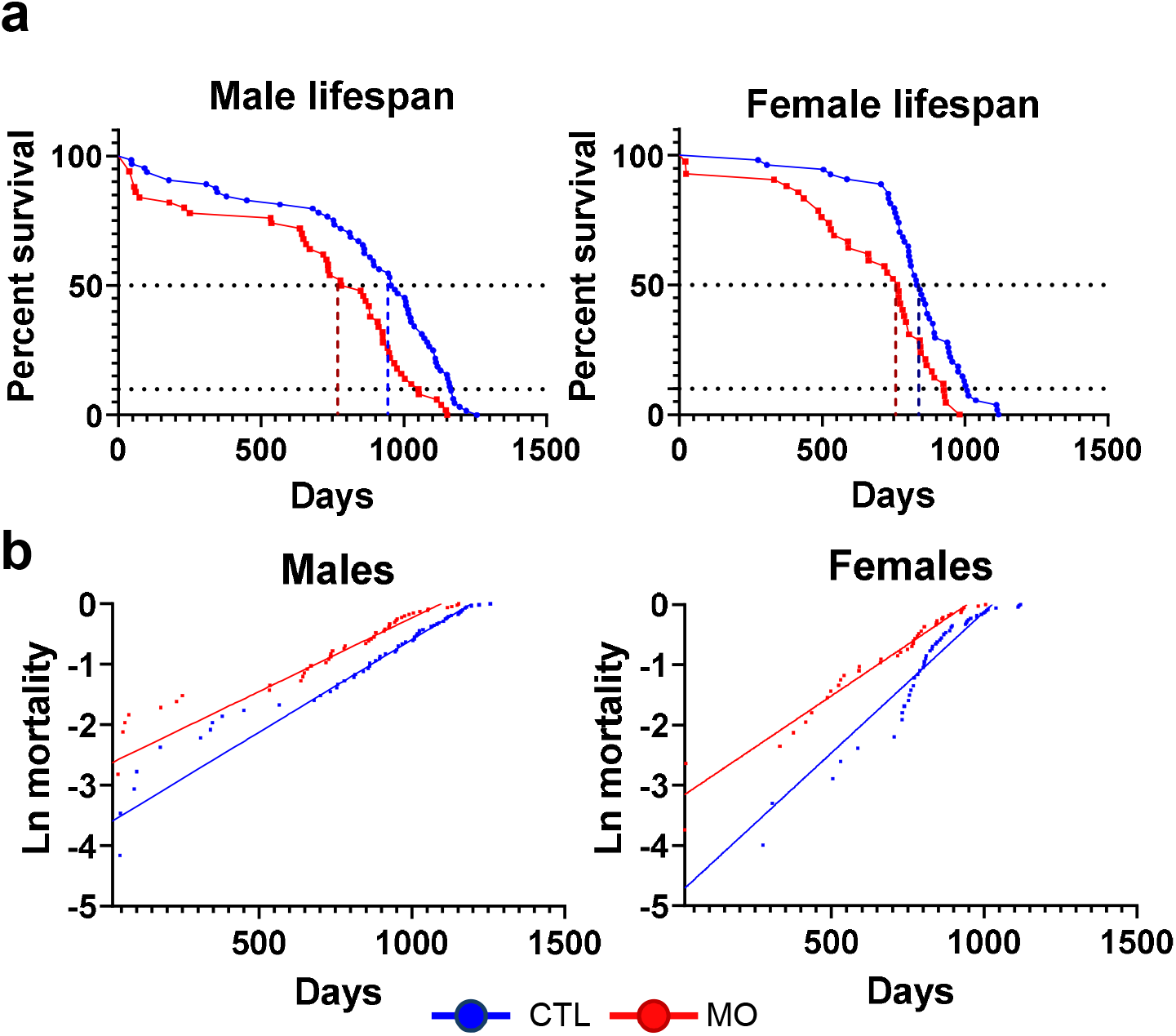
Maternal obesity reduces the lifespan of the offspring. The survival data are shown for the male and female offspring described in Fig. S1A born to female dams fed control diet (Ctl, blue) or western diet (maternal obesity, red). **A** shows the Kaplan-Meier survival curves with the dashed lines indicating the 50^th^ and 90^th^ percentile survival. The number of mice in each group and the analysis of the survival data are given in Table 1 showing that maternal obesity significantly reduced the lifespan of both male (p < 0.01) and female (p < 0.001) mice compared to control mice. **B** Gompertz plots of survival data are shown using the distributions of ages at death from the equation u_x_ = ae^bx^, where u_x_ is the instantaneous age-specific mortality. The Gompertz curves were significantly different (p<0.0001) for both male and female maternal obesity mice compared to control. The initial rate of mortality and the rate of increase in the hazard for mortality (i.e., the mortality rate) are given in Table 1.

To gain insight into how maternal obesity reduced the lifespan of the offspring, we performed a Gompertz mortality analysis on the lifespan data which can be used to compare mortality patterns between groups within a species^20,21^. Gompertz analysis of any species shows that the adult mortality/aging rate increases exponentially with age indicating that probability of death at any given time is much higher at older ages than at younger ages^20^. As explained by Kirkwood, 2015, the μ(x) in the gompertz equation (μ(x) = αe^βx^) denotes the age-specific mortality (mortality rate at age x). The β denotes the overall rate of mortality/aging i.e., ‘the time required for the death rate to increase by a factor of two’, and the ‘y’ intercept α denotes the initial mortality rate which essentially measures the initial vulnerability to disease before the onset of aging^20^. As shown in Fig. 1B and Table 1, the Gompertz curves were overall significantly different (p<0.0001) for both male and female maternal obesity offspring compared to control offspring. Unexpectedly, we found that the ‘β’ value (the overall rate of mortality/aging) was reduced by 27 and 20% in female and male maternal obesity offspring, respectively, compared with control mice, suggesting that the reduced lifespan of the offspring born to obese dams was not due to an increase in rate of aging. The y-intercept of the gompertz regression shows that the initial mortality (α) rate, which occurs well before the onset of aging, is dramatically increased by 161% in the male offspring and >387% in the female offspring. Therefore, the increased age-specific mortality [μ(x)], observed in the maternal obesity offspring is due to the increased initial death rate that occurs before onset of aging and remained high throughout lifespan of the male and female offspring. In addition to the Gompertz analysis, we tested for the proportional hazards assumption in the mixed effects Cox model. We determined that the assumption holds between maternal obesity and control offspring for both males (p < 0.95) and females (p < 0.2), meaning that the hazard ratio does not appreciably change with respect to age. These data support our findings from the Gompertz analysis that increased mortality in maternal obesity is driven by age-independent factors, or “starting” biological age which results in higher mortality throughout lifespan instead of factors that modify the rate of aging.

### 2.3 Offspring Fibroaging

An age-related increase in pathology leading to increased incidence of disease is a hallmark of aging. Fibrosis (fibroaging) is recognized as a marker of aging pathology^22^, which involves an excessive age-related accumulation of fibrous connective tissue (e.g., collagen) in most tissues^23^. In addition, an increase in markers of fibrosis has been reported in embryo and fetal tissue from mice^24^ and non-human primates^25^ born to obese mothers. Therefore, we measured the level of fibrosis in three tissues from 25-month-old control and maternal obesity offspring. Using either picrosirius red (PSR) or Masson’s trichrome (MT) staining, we first quantified the level of collagen deposition in kidney, liver, and heart of male and female control and maternal obesity offspring (Fig. 2A, B). Collagen deposition in kidney occurred around renal tubules and blood vessels and was significantly increased (3-to 4-fold) in both male and female maternal obesity offspring compared to control mice. We also quantified the level of collagen type I fibers in the collagen deposits because fibrosis involves an increase in thick collagen type I fibers^26^. A ∼2-fold increase in type I fibers was observed in maternal obesity male and female offspring. The liver also showed a significant increase (2-to 3-fold) in collagen deposition in both male and female maternal obesity o that occurred around the portal triads and was associated with a ∼2-fold increase in type I fibers. Similar results were found in the maternal obesity offspring heart with a significant increase (∼2-fold) in collagen deposition around the perivascular region and interstitial spaces and over a 2-fold increase in type I fibers in the collagen deposits.

**Fig. 2:**
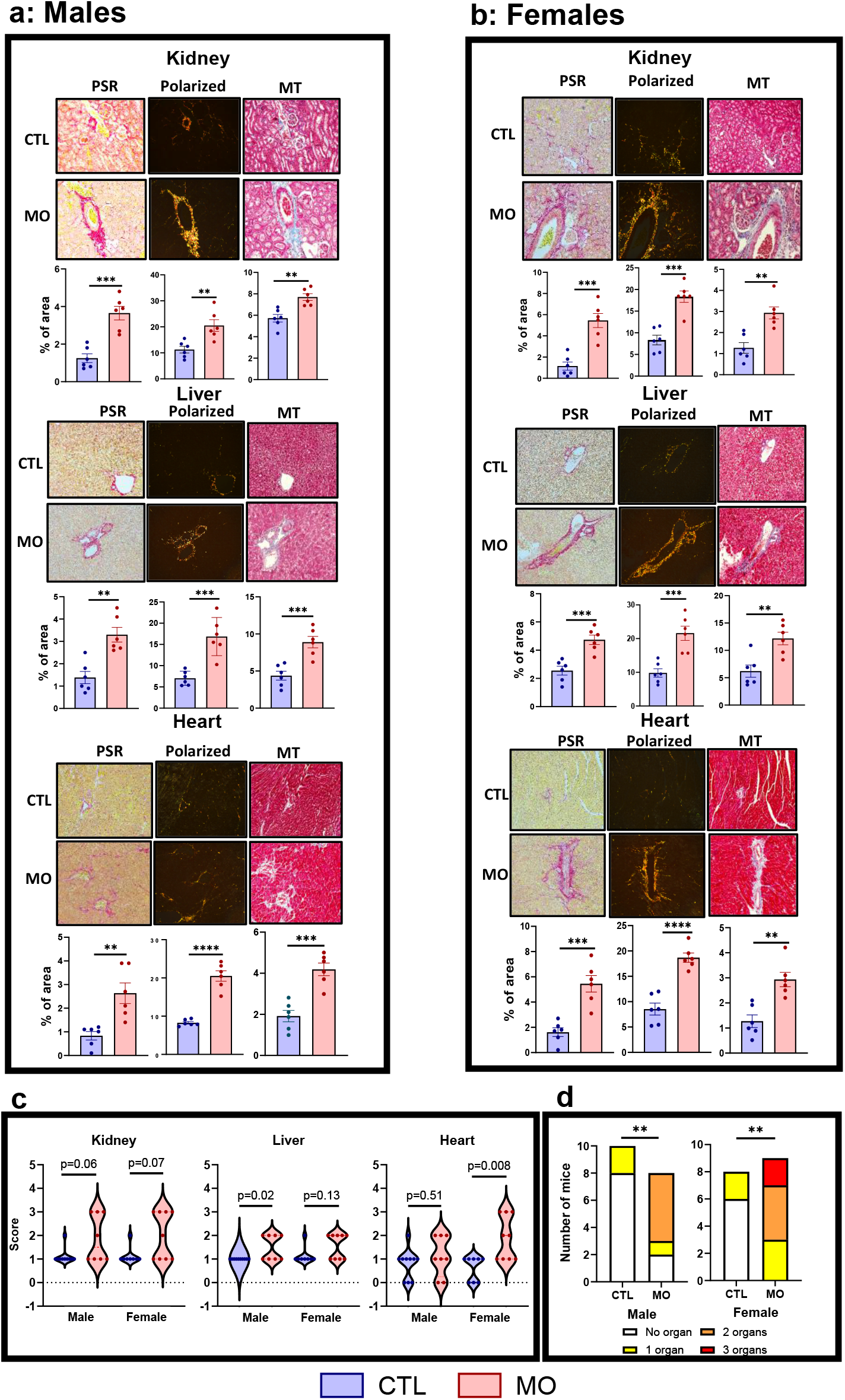
Maternal obesity promotes fibrosis in offspring. Fibrosis was measured in the kidney, liver, and heart of 26-month-old male (**A**) and female (**B**) mice born to either control (Ctl; blue bars) or maternal obese (maternal obesity; red bars) dams. **A & B** show collagen deposition in the three tissues using picrosirius red (PSR) and Masson’s trichome (MT) staining as previously described^54^. The left column shows representative images of PSR staining (20x) showing collagen deposition (red) in the periglomerular, tubulo-interstitial regions, and perivascular regions of the kidneys; the portal triad of the liver; and the sub-endocardial and transmural regions of the heart. The middle columns (Polarized) display PSR-stained sections under polarized light (20x) showing yellow-orange birefringence of thick, mature collagen type I fibers. The right columns show representative MT staining (20x), highlighting collagen deposition (blue) in the same anatomical regions as with PSR staining. Quantification of the collagen-positive area per field was measured as we have described^54^ and is shown in the graphs below each image with data represented as mean ± SEM for 6 mice with individual data points shown. Statistical comparisons between the control and maternal obesity groups were performed using a two-tailed t-test (*p < 0.05, **p < 0.01). **C** shows the fibrosis scoring liver^38^, kidneys^39^, and heart^40^, which was performed by three veterinary pathologists blinded to treatment groups. Data are presented as the mean ± SEM for 8 to 10 mice with individual data points shown. Statistical comparison p values are given for the fibrosis scores as measured by the Mann-Whitney U test. **D** shows the number of mice with mild to moderate fibrosis (fibrosis scores of 2 or 3) in one or more tissues of each mouse. Statistical comparisons between the control and maternal obesity groups for each sex were performed using Fisher’s exact test. (**p < 0.01).

We also assessed the severity of fibrosis in the three tissues using pathology scoring systems based on the location and extent of fibrosis in the three tissues that ranged from 0 (no fibrosis) to 4 (severe fibrosis). As shown in Supplementary Table 1, the severity of fibrosis in the three tissues from the 18 control male and female offspring ranged from 0 to 1 (few collagen fibers) for 14 of the mice with only 4 showing a score of 2 (mild fibrosis) in one of the three tissues. In contrast, 16 of the 17 male and female maternal obesity offspring had a score of 2 or 3 (moderate fibrosis) in one or more tissues. As shown in Fig. 2C, the mean severity of fibrosis was greater for both male and female offspring from maternal obesity mice in all three tissues with the increase significant for the liver in males and for the heart in females. The data in Fig. 2D and Supplementary Table 1 show the number of mice that have mild to moderate fibrosis (i.e., scores of 2 or 3) in the three tissues from each mouse. We found that over 50% of the male and female maternal obesity offspring showed mild to moderate fibrosis in multiple tissues in the same mouse compared to none for the 18 control mice, i.e., none of the control offspring showed mild to moderate fibrosis in more than 1 tissue. In female maternal obesity offspring, 2 of the 9 mice showed mild to moderate fibrosis in all three tissues. Because the increase in fibrosis with age has been linked to organ dysfunction and has been shown to contribute to a variety of age-related diseases^22, 27, 28^, our data suggest that old maternal obesity offspring would likely show an increase in multiple organ dysfunction and incidence of age-related diseases. For example, increased cardiac fibrosis found in maternal obesity offspring is a strong biomarker, particularly for arrhythmias, heart failure, and increased cardiovascular mortality^29^. Likewise, increased fibrosis observed in the kidneys of maternal obesity mice would affect the glomeruli, tubule-interstitial space and arterial vessel, which are indicators of chronic kidney disease^30^ and could possibly contribute to the early death observed in the maternal obesity offspring.

## 3. Discussion

Our study clearly demonstrates that maternal obesity has a long-term negative impact on both male and female offspring as evidenced by reduced lifespan and an increase in fibrosis in tissues from old mice. These data suggest that maternal obesity acts as a prenatal/perinatal insult that significantly impacts multiple tissues in these offspring to decrease longevity. Importantly, the changes occurred in the maternal obesity offspring even though they were maintained on a non-obesogenic control diet, did not experience excess body weight gain, and had minimal environmental stress throughout their lives. While several studies have shown that pre-gestational and gestational maternal obesity differentially programs the offspring, we chose to follow the continuous period of western diet exposure from pre-pregnancy to lactation in combination with maternal obesity to maximally affect the offspring and to more closely resemble the human conditions of pregnancy and lactation. Having established that life-span is indeed shortened, and that the early pre-peri-fetal and post-natal effects impact lifespan, our future studies will focus on parsing out the effects in early life. Our Gompertz data provides an insight into how maternal obesity might impact aging and lifespan. The reduced lifespan in the maternal obesity offspring was not due to an increase in the overall rate of mortality/aging. Rather, the initial mortality/death rate, which provides the starting point of the exponential increase in mortality with age^31^, was elevated in the maternal obesity offspring resulting in a higher mortality at all ages. Our data indicates that the long-term effects of maternal obesity on lifespan and pathology are primarily due to early effects of maternal obesity on the developing fetus/neonate, long before the emergence of disease. These results are consistent with studies in human and non-human primates demonstrating early epigenetic changes^32, 33^ due to maternal obesity/poor diet which in turn may predispose offspring to metabolic, cardiovascular and other non-communicable inflammatory diseases later in life including accelerated epigenetic aging^34^. These results also suggest that early-life interventions are likely to alleviate the long-term adverse effects of maternal obesity on the offspring. Importantly, interventions would not have to be employed throughout the life of the maternal obesity offspring because maternal obesity did not accelerate the mortality rate. Several studies have reported that early interventions can attenuate adverse effects of maternal obesity observed in adult offspring as reviewed by Shrestha et al^35^. For example, exercise during pregnancy has been shown to improve insulin resistance^36^, cardiovascular disease^37^, and reduce metabolic dysfunction-associated fatty liver disease ^38^ in adult mice born to obese dams. In addition, anti-oxidants or anti-inflammatory agents such as pyrroloquinoline quinone^39^, N-acetylcysteine^40^, and Omega 3-fatty acids^41^, given during pregnancy and lactation to mothers fed a western style diet resulted in adult offspring with improved glucose tolerance and reduced steatosis^42^ as well as reduced liver fibrosis^43,44^. Based on our Gompertz data, we predict that these interventions might have a pronounced effect on longevity. Also, it is important to understand the underlying mechanism(s) driving the negative effects of maternal obesity which will also help in the development of interventions that could prevent or alleviate the negative effects of maternal obesity on the offspring. Various pathways associated with aging has been shown to be dysregulated in MO offspring e.g., oxidative stress, microbiome changes, DNA damage, inflammation, mTOR activation and mitochondrial dysfunction^45^. Therefore, further studies identifying mechanisms, interventions and the window of treatment that can counter the negative effects of maternal obesity have the potential of allowing children born to the growing number of mothers with obesity to live out their lifespan with improved healthspan and lifespan.

### 3.1 Limitations

One of the limitations of the study is that the pathological data are limited to fibrosis (fibroaging) at 26-months of age. More extensive pathology analysis of the mice and at earlier ages as well as at the end-of-life pathology when the mice die would show if pathological lesions occur earlier in the MO offspring and if the MO and control offspring are dying from similar or different pathologies. The other limitation of our study is the lack of information about the healthspan (e.g., frailty, physical activity, cognition etc.) of the aged maternal obesity offspring. While our current study provides strong evidence that lifespan is reduced and fibroaging is increased in tissues of the maternal obesity offspring, it would be important to show how maternal obesity impacts the healthspan of the offspring. And although, the detrimental effects of maternal obesity on developmental programming on the fetus have been well documented in mice, monkey, and human pregnancy^41, 46-51^ our study also lacks the early-life phenotypic data, which could have provided additional validation for the Gompertz data. Finally, while all studies on maternal obesity have used inbred C57BL/6 mice as we have, it would be translationally relevant to show that the impact of maternal obesity on lifespan and aging is observed also on a genetic diverse background, e.g., UM-HET3 mice used by the Intervention Testing Program (National Institute of Aging) to study the impact of compounds on the lifespan of mice.

## 4. Materials and methods

### 4.1 Animal model

Male and Female C57BL/6J mice were obtained from Jackson Laboratories (Bar Harbor, Maine, USA). All animals were generated and maintained at the University of Oklahoma Health Campus under SPF conditions in a HEPA barrier environment. They were housed in ventilated cages at 20□±□2°C with controlled 12-hour light/dark cycles and fed *ad libitum*. All females used to generate offspring were tested for fertility (mated and had first litter) at 6 weeks of age. Only females found to have litters were randomly assigned to either control diet (CD, PicoLab Rodent Diet 20) or western diet (WD, Teklad TD.8813) feeding (See Supplemental Fig. 1 for diet composition) for two months based on previously established regimens^42, 52, 53^. We had 12 females on CD and 19 females on WD. After two months of feeding, body composition was analyzed to ensure that western diet feeding induced obesity. Both control diet and western diet groups were then mated with control diet fed males and continued with their respective diets during pregnancy and lactation period. Continuous WD exposure was employed to create an obesogenic environment that closely resembles the chronic obesogenic environment found in humans. We paired the male to female overnight and on observation of vaginal plug next morning the males are removed. This strategy was followed to ensure that males get less than 16-hours of WD exposure. If vaginal plugs were not observed next morning, the male was removed and new male introduced to the breeding cage. Details of the litter size and number of litters per dam are given in supplementary Table 1. The litters were left with their respective mothers during the lactation period without any litter standardization. The resulting pups (control and maternal obesity groups) were weaned on to a non-obesogenic control diet regardless of maternal diet at 21 days of age, group housed (4-5 mice per cage) and allowed to age without any further dietary intervention or stress. Mice were euthanized using isoflurane and exsanguination, and tissues were harvested for pathology. Control diet and maternal obesity offspring were not exposed to any manipulations other than cage changes every other week. Offspring were monitored daily for health and morbidity and allowed to die naturally unless meeting near-death humane conditions (Inability to move to access food and water, large tumors, major weight loss of >20%) where mice typically die within 24-48 hours. All animal experiments were performed according to protocols approved by the Institutional Animal Care and Use Committee.

### 4.2 Body composition

Body composition was determined using nuclear magnetic resonance spectroscopy (NMR Bruker minispec). Body weight, fat mass, lean mass, and fluid mass were recorded in grams in the mothers prior to breeding to ensure obesity in western diet fed females. Body composition was obtained longitudinally at ages of 3 months, 6 months, 12 months, 18 months, and 24 months in the male and female maternal obesity offspring.

### 4.3 Fibrosis scoring

Formalin-fixed liver, kidneys, and heart were embedded in paraffin, and 4 µm serial sections were generated using a microtome. Serial sections were then either used for either picrosirius red (PSR) or Masson’s trichrome (MT) staining as previously described.^37^ For staining, sections were deparaffinized and stained with either PSR or MT for 1 hour. Excess stain was removed by rinsing with acidified water. Sections were then dehydrated with ethanol and cleared with xylene. Whole sections were then imaged using a Nikon TI Eclipse microscope (Nikon, Melville, NY) at 20x magnification. Staining quantification of collagen-positive area per field was measured as average percent of area showing staining using Image J software. Fibrosis scoring was performed according to the following scoring systems: Liver: Brunt scoring criteria^38^: 0 = none, 1 = mild perisinusoidal fibrosis, 2 = zone 3 periportal fibrosis, 3 = bridging fibrosis, and 4 = cirrhosis; Kidney: The GTIA regions of the kidneys [i.e., Glomeruli(G), tubulointerstitial (TI), and arterial vessels(A)] were scored as follows^39^: 0 = none, 1 = few collagen fibers (0-20% affected), 2 = mild fibrosis (21-40% affected), 3 = moderate (41-60% affected), and 4 = severe (61-80% affected); Heart^40^: 0 = none, 1 = a few scattered collagen fibers throughout the heart, 2 = patchy fibrosis in 1 to 5 areas (<20% of the field), contiguous subendocardial fibrosis involving less than half the circumference, or transmural fibrosis involving less than half the circumference, 3 = patchy fibrosis in more than 5 areas (> 20% of the field) with contiguous subendocardial fibrosis involving more than half the circumference or transmural fibrosis of more than half the circumference, and 4 = total transmural fibrosis.

### 4.4 Statistics

Graphs and statistical analysis were performed using GraphPad Prism v 10.6 (GraphPad Software, Boston, Massachusetts USA). Analysis for mixed effects Cox proportional hazards model, linear mixed models, quantile mixed regression, and Gompertz mortality analysis was performed in SAS software version 9.4 (SAS Institute, Cary, North Carolina, USA). Body compositional data from the mothers was compared using two-tailed Welch’s t-tests while longitudinal changes in the offspring were compared with 2-way ANOVA or mixed model if data was missing due to deaths. We utilized a Log-rank (Mantel-Cox) test and a Gehan-Breslow-Wilcoxon test to compare the Kaplan-Meier survival data in the lifespan cohort. To compare mean, median, and 90^th^ percentile survival time, we used a linear mixed model and a quantile mixed regression, respectively. To control for cage effects, we fit a mixed effects Cox proportional hazards model with cage as a random effect and checked the proportional hazards assumption. Gompertz mortality analysis was conducted starting with a log-linear transform of mortality data, which was then used to fit a least-squares linear regression. The y-intercept corresponds to the initial mortality parameter α of the Gompertz equation, while the slope corresponds to the aging rate parameter β. Type 3 SS ANOVA was used to test for significant differences in Gompertz curve parameters. The original form of the Gompertz equation is

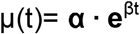

To compare maximum lifespans, we employed the Max lifespan test developed by Gao et al.^19^ Comparisons for fibrosis area were made with two-tailed student’s t-test, and fibrosis scoring was compared by Mann-Whitney U test. The number of mice with mild to moderate fibrosis scores (fibrosis score >= 2) in one or more tissues of each mouse were compared using a Fisher’s exact test.

## Figure Legends and Tables

Data are given in days with the percent difference in the maternal obesity mice from the control mice shown in parentheses. ^a^Linear Mixed Models used to test significance difference in mean lifespan; ^b^Quantile mixed regression used to test the significant difference in median and 90^th^ percentile lifespans; ^c^Maximum lifespan test developed by Gao et al^19^ to test for significant differences in the upper tail of the distribution in the survival data; ^d^Age when the last mouse in the group died; ^e^Cox Proportional hazards assumption holds for maternal diet. Model controls for random cage effects. ^f^Mixed effects Cox proportional hazards analysis and ^g^Gehan-Breslow-Wilcoxon proportional hazards models with Gompertz distribution used to test for significant differences in lifespan curves between the control and maternal obesity mice for each sex; ^h^The Gompertz a (initial rate of mortality) and b (mortality rate) were calculated from the data in Figure 1B using the distributions of ages at death by the equation u_x_ = ae^bx^, where u_x_ is the instantaneous age-specific mortality. ^i^Type 3 SS ANOVA was used to test for significant difference in the Gompertz curve parameters.

## Supplemental figures and tables

**Supplemental Fig. 1: Experimental design. A** shows breeding scheme for the maternal obesity (maternal obesity) model studied. The mice were generated and maintained in the animal facility at The University of Oklahoma Health Campus. Female C57BL/6J mice obtained from the Jackson Laboratory were mated once at 2 months of age to test fertility. Females were then placed on either a control diet (control diet, PicoLab Rodent Diet 20) or western diet (western diet, Teklad TD.88137) for 2-months. Body composition was then performed to confirm obesity in maternal obesity fed females. Females from both groups were then mated to control diet-fed male C57BL/6J mice, and their respective diets were continued through pregnancy and lactation. The resulting pups were then weaned onto control diet and group housed (4 to 5 mice/cage) in ventilated cages at 20□±□2°C, on a12-h/12-h dark/light cycle. All procedures were approved by the Institutional Animal Care and Use Committee at The University of Oklahoma Health Campus. **B** shows the macronutrient composition of the control diet and western diet diets. **C** shows the body composition of dams fed control diet (blue bars) or western diet (red bars). Fat mass and lean mass were determined by nuclear magnetic resonance spectroscopy (NMR Bruker minispec). The data (5 mice per group) are expressed as the mean ± SEM, and the individual data points shown. Statistical significance was determined using two-tailed Welch’s t-test (* p < 0.05; **** p < 0.0001).

**Supplemental Fig. 2: Body composition of control and maternal obesity mice.** The longitudinal body weights and composition determined by NMR Bruker minispec) were measured at 3, 12, 18, and 24 months of age for each male (**A**) and female (**B**) control (blue) and maternal obesity (red) mouse. The data are expressed as the mean ± SEM for 9-12 per mice group. Using a two-way ANOVA (or mixed model if data were missing due to deaths), no statistically significant difference was found between the maternal obesity and control mice for either male or female mice for any of the measurements of body composition.

**Supplemental Fig. 3: Whole section fibrosis images for heart, kidney, and liver.** Whole sections of kidney, liver, and heart of 26-month-old male (**A**) and female (**B**) mice born to either control (Ctl) or maternal obese (maternal obesity) dams. **A & B** show collagen deposition in the three tissues using picrosirius red (PSR) and Masson’s trichome (MT) staining as previously described^54^. The left column shows representative images of PSR staining showing collagen deposition (red) in the kidneys, liver, and heart. The right columns show representative MT staining, highlighting collagen deposition (blue) in sections serial to the ones used in PSR staining.

**Supplementary Table 1.** List of number of Litters/dam and Litter Size.

| Control Diet |  |  |  | Lifespan |  | Body Composition |  | Fibrosis |  |
| --- | --- | --- | --- | --- | --- | --- | --- | --- | --- |
| Dam ID | Dam # | Litter # | Litter Size | # of Male | # of Female | # of Male | # of Female | # of Male | # of Female |
| 60697 | 1 | 1 | 10 | 5 | 5 | 0 | 0 | 0 | 0 |
|  |  | 2 | 3 | 3 | 0 | 0 | 0 | 0 | 0 |
|  |  | 3 | 9 | 0 | 1 | 0 | 0 | 0 | 0 |
| 60698 | 2 | 1 | 2 | 2 | 0 | 0 | 0 | 0 | 0 |
|  |  | 2 | 8 | 1 | 3 | 0 | 0 | 3 | 2 |
| 69028 | 3 | 1 | 8 | 2 | 5 | 0 | 5 | 0 | 0 |
|  |  | 2 | 9 | 2 | 3 | 0 | 0 | 1 | 2 |
| 69056 | 4 | 1 | 7 | 2 | 1 | 0 | 0 | 0 | 0 |
|  |  | 2 | 5 | 0 | 2 | 0 | 0 | 0 | 2 |
|  |  | 3 | 9 | 3 | 0 | 0 | 0 | 0 | 0 |
| 69922 | 5 | 1 | 6 | 2 | 4 | 0 | 0 | 0 | 0 |
| 69940 | 6 | 1 | 9 | 3 | 0 | 0 | 0 | 0 | 0 |
|  |  | 2 | 7 | 3 | 0 | 0 | 0 | 0 | 0 |
| 69941 | 7 | 1 | 10 | 8 | 2 | 0 | 0 | 0 | 0 |
|  |  | 2 | 10 | 0 | 6 | 0 | 0 | 0 | 1 |
| 69946 | 8 | 1 | 8 | 5 | 3 | 3 | 3 | 0 | 0 |
|  |  | 2 | 2 | 2 | 0 | 0 | 0 | 0 | 0 |
|  |  | 3 | 5 | 5 | 0 | 0 | 0 | 0 | 0 |
| 69947 | 9 | 1 | 7 | 2 | 5 | 0 | 0 | 0 | 0 |
|  |  | 2 | 2 | 0 | 0 | 0 | 0 | 2 | 0 |
| 69948 | 10 | 1 | 7 | 2 | 4 | 0 | 0 | 0 | 0 |
|  |  | 2 | 9 | 4 | 1 | 0 | 0 | 2 | 1 |
| 69949 | 11 | 1 | 10 | 5 | 4 | 5 | 4 | 0 | 0 |
|  |  | 2 | 2 | 0 | 0 | 0 | 0 | 2 | 0 |
| 69968 | 12 | 1 | 8 | 3 | 5 | 2 | 0 | 0 | 0 |
|  |  |  | Total | 64 | 54 | 10 | 12 | 10 | 8 |

| Western Diet |  |  |  | Lifespan |  | Body Composition |  | Fibrosis |  |
| --- | --- | --- | --- | --- | --- | --- | --- | --- | --- |
| Dam ID | Dam # | Litter # | Litter Size | # of Male | # of Female | # of Male | # of Female | # of Male | # of Female |
| 2230 | 1 | 1 | 4 | 2 | 0 | 0 | 0 | 0 | 0 |
|  |  | 2 | 3 | 3 | 0 | 0 | 0 | 0 | 0 |
| 2292 | 2 | 1 | 4 | 3 | 0 | 0 | 0 | 0 | 0 |
| 2297 | 3 | 1 | 5 | 0 | 5 | 0 | 0 | 0 | 0 |
|  |  | 2 | 4 | 2 | 0 | 0 | 0 | 0 | 0 |
| 2298 | 4 | 1 | 8 | 3 | 4 | 0 | 0 | 0 | 0 |
| 69027 | 5 | 1 | 3 | 2 | 0 | 0 | 0 | 0 | 0 |
| 69053 | 6 | 1 | 2 | 1 | 1 | 0 | 0 | 0 | 0 |
| 69062 | 7 | 1 | 9 | 3 | 0 | 3 | 0 | 0 | 0 |
|  |  | 2 | 4 | 3 | 0 | 0 | 0 | 0 | 0 |
| 69068 | 8 | 1 | 10 | 5 | 5 | 5 | 5 | 0 | 0 |
|  |  | 2 | 6 | 2 | 1 | 2 | 0 | 0 | 0 |
| 69082 | 9 | 1 | 3 | 3 | 0 | 0 | 0 | 0 | 0 |
|  |  | 2 | 9 | 3 | 1 | 0 | 0 | 2 | 3 |
| 69084 | 10 | 1 | 9 | 4 | 1 | 0 | 0 | 0 | 0 |
| 69091 | 11 | 1 | 3 | 3 | 0 | 0 | 0 | 0 | 0 |
|  |  | 2 | 2 | 2 | 0 | 0 | 0 | 0 | 0 |
| 69938 | 12 | 1 | 8 | 4 | 4 | 0 | 0 | 0 | 0 |
|  |  | 2 | 4 | 2 | 2 | 0 | 0 | 0 | 0 |
| 69979 | 13 | 1 | 3 | 3 | 0 | 0 | 0 | 0 | 0 |
|  |  | 2 | 5 | 0 | 0 | 0 | 0 | 2 | 0 |
| 69050 | 14 | 1 | 2 | 0 | 2 | 0 | 0 | 0 | 0 |
| 69086 | 15 | 1 | 5 | 0 | 3 | 0 | 0 | 0 | 2 |
|  |  | 2 | 7 | 0 | 5 | 0 | 5 | 2 | 0 |
| 69090 | 16 | 1 | 2 | 0 | 2 | 0 | 0 | 0 | 0 |
|  |  | 2 | 2 | 0 | 1 | 0 | 0 | 0 | 1 |
| 69092 | 17 | 1 | 4 | 0 | 2 | 0 | 0 | 0 | 2 |
| 69961 | 18 | 1 | 5 | 0 | 3 | 0 | 0 | 0 | 1 |
| 69057 | 19 | 1 | 2 | 0 | 0 | 0 | 0 | 2 | 0 |
|  |  |  | Total | 53 | 42 | 10 | 10 | 8 | 9 |

Number of litters and litter size for each dam, as well as sex and experiment distribution from each litter are shown.

**Supplementary Table 2.** Fibrosis scores for 24-month-old control and maternal obesity mice.

|  | Liver | Heart | Kidney |  | Liver | Heart | Kidney |
| --- | --- | --- | --- | --- | --- | --- | --- |
| **Control Male Mice** |  |  |  | **Control Female Mice** |  |  |  |
| Mouse 1 | 1 | 2 | 1 | Mouse 19 | 1 | 1 | 1 |
| Mouse 2 | 1 | 1 | 1 | Mouse 20 | 1 | 1 | 1 |
| Mouse 3 | 1 | 1 | 1 | Mouse 21 | 1 | 0 | 1 |
| Mouse 4 | 1 | 1 | 1 | Mouse 22 | 2 | 1 | 1 |
| Mouse 5 | 1 | 0 | 1 | Mouse 23 | 1 | 1 | 1 |
| Mouse 6 | 1 | 0 | 2 | Mouse 24 | 1 | 0 | 2 |
| Mouse 7 | 1 | 0 | 1 | Mouse 25 | 1 | 0 | 1 |
| Mouse 8 | 1 | 1 | 1 | Mouse 26 | 1 | 1 | 1 |
| Mouse 9 | 1 | 1 | 1 |  |  |  |  |
| Mouse 10 | 1 | 1 | 1 |  |  |  |  |
| **Maternal Obesity Male Mice** |  |  |  | **Maternal Obesity Female Mice** |  |  |  |
| Mouse 11 | 2 | 2 | 1 | Mouse 27 | 2 | 1 | 1 |
| Mouse 12 | 1 | 2 | 1 | Mouse 28 | 1 | 1 | 3 |
| Mouse 13 | 2 | 1 | 1 | Mouse 29 | 2 | 3 | 1 |
| Mouse 14 | 2 | 1 | 3 | Mouse 30 | 1 | 1 | 3 |
| Mouse 15 | 1 | 2 | 3 | Mouse 31 | 2 | 2 | 2 |
| Mouse 16 | 2 | 1 | 3 | Mouse 32 | 2 | 2 | 1 |
| Mouse 17 | 1 | 0 | 2 | Mouse 33 | 2 | 3 | 3 |
| Mouse 18 | 1 | 0 | 1 | Mouse 34 | 1 | 1 | 3 |
|  |  |  |  | Mouse 35 | 1 | 3 | 1 |

Fibrosis scores for each tissue for each mouse are shown. The scores were determined by three pathologists independently as a double-blind study using picrosirius red staining at 20x magnification with a total of 5-6 slides per group. Slides for liver^55^, kidneys^56^, and heart^57^ were then scored for fibrosis using established scoring systems as described in methods

