## Supplemental figures for "Maternal Obesity Decreases Offspring Lifespan"

### Slide 1
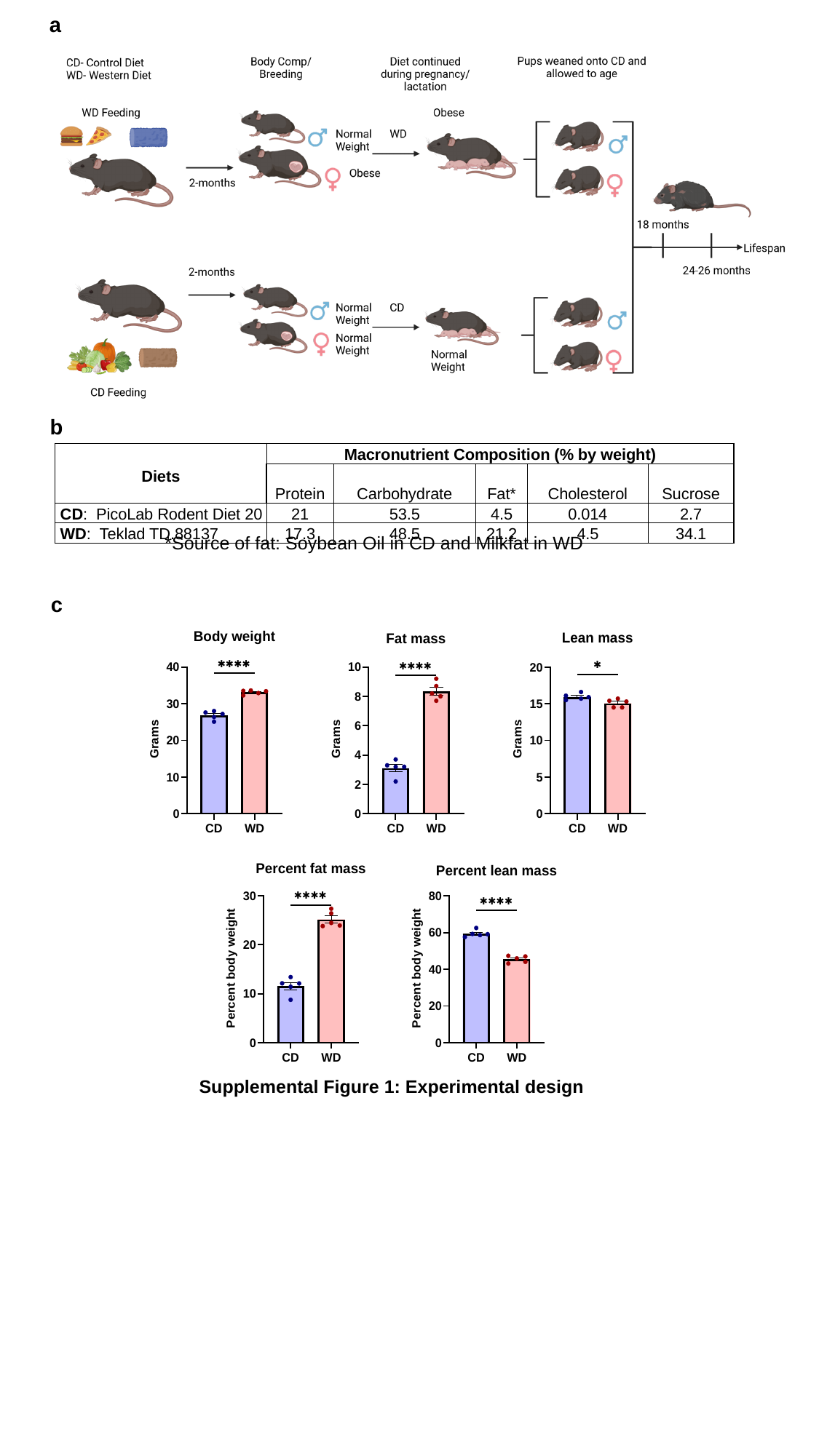

a
b
| Diets | Macronutrient Composition (% by weight) | | | | |
| --- | --- | --- | --- | --- | --- |
| | Protein | Carbohydrate | Fat\* | Cholesterol | Sucrose |
| CD: PicoLab Rodent Diet 20 | 21 | 53.5 | 4.5 | 0.014 | 2.7 |
| WD: Teklad TD.88137 | 17.3 | 48.5 | 21.2 | 4.5 | 34.1 |
*Source of fat: Soybean Oil in CD and Milkfat in WD
c
Supplemental Figure 1: Experimental design

### Slide 2
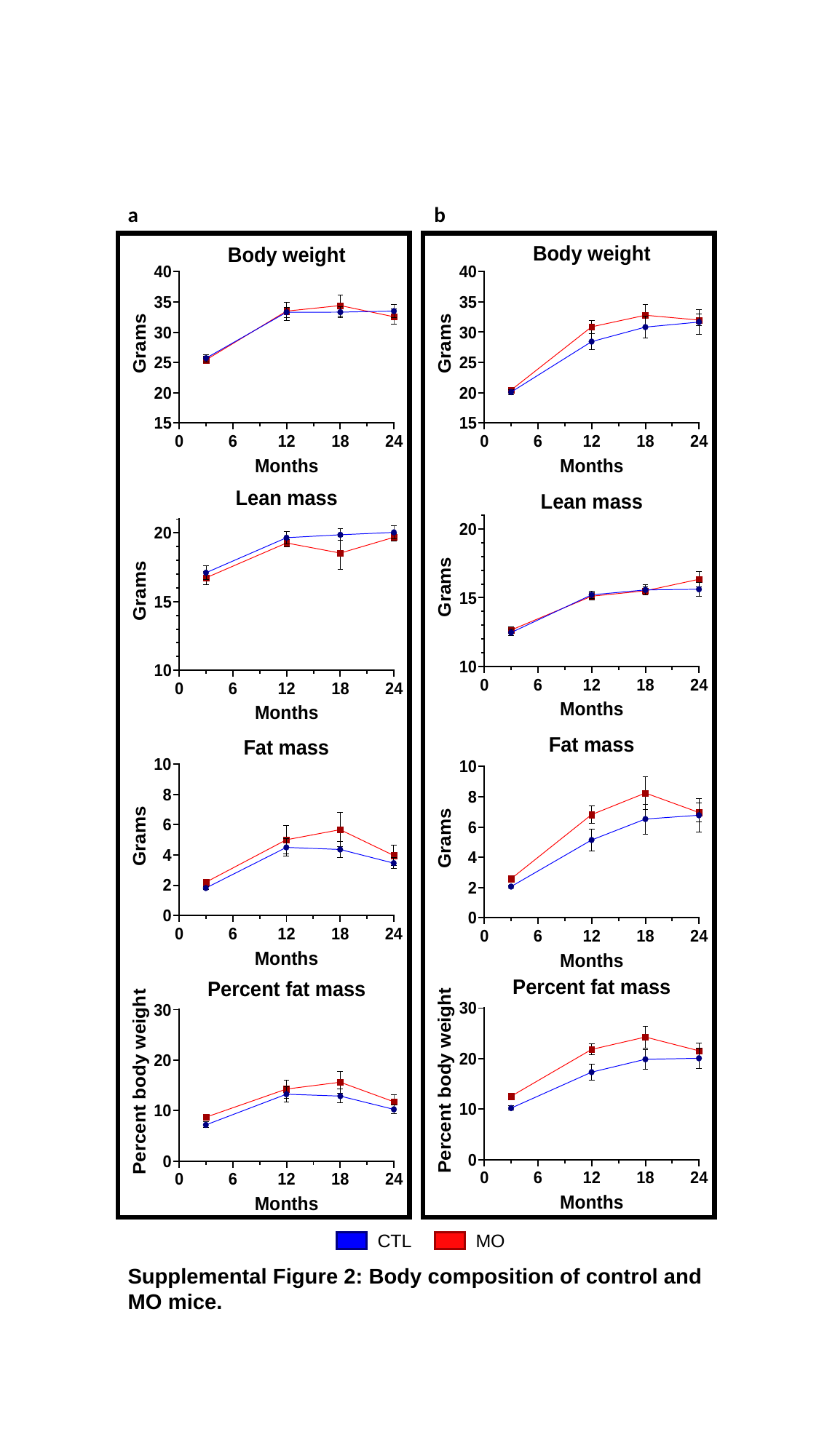

a
b
CTL
MO
Supplemental Figure 2: Body composition of control and MO mice.

### Slide 3
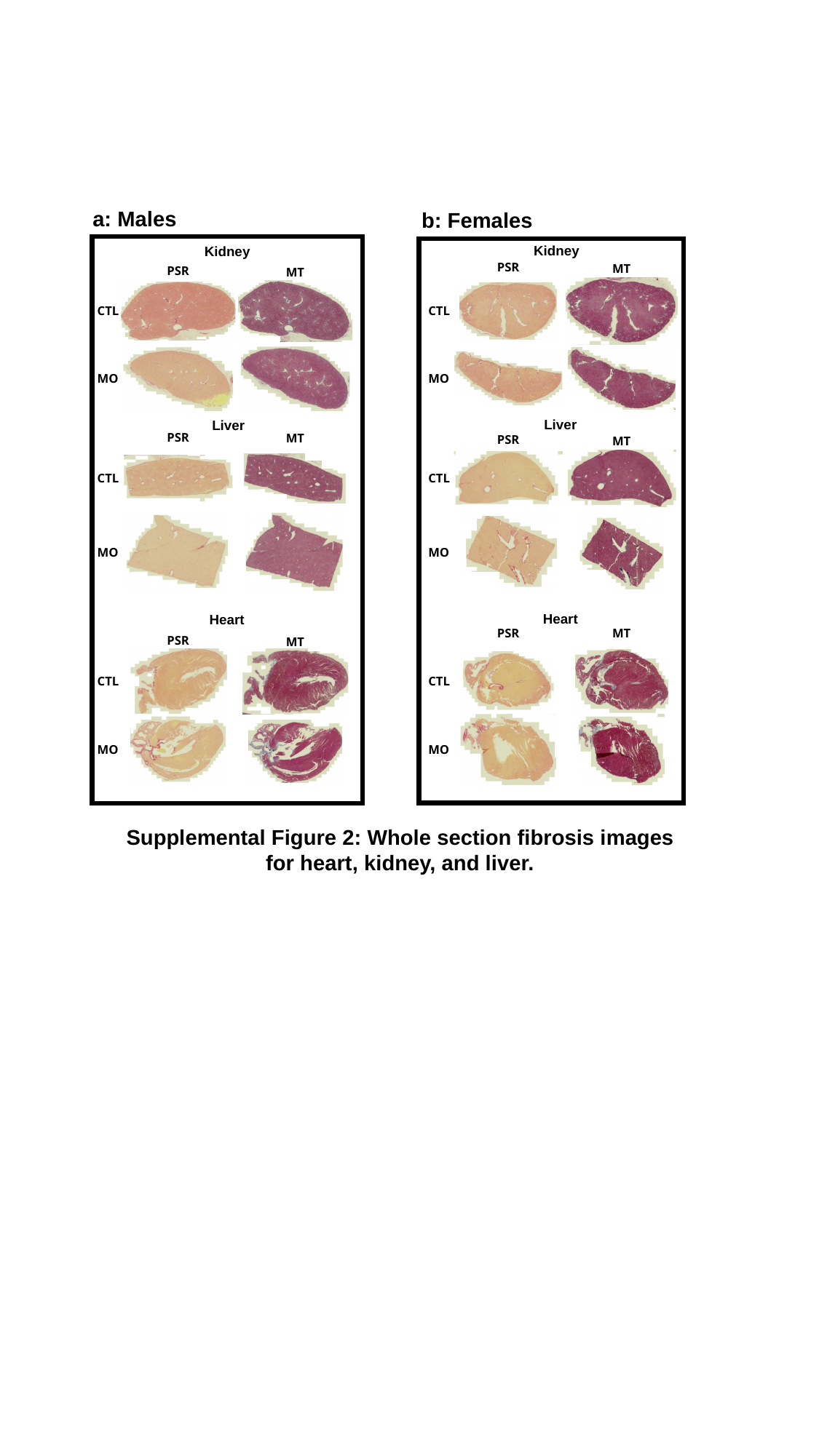

a: Males
b: Females
Kidney
Kidney
PSR
MT
PSR
MT
CTL
CTL
MO
MO
Liver
Liver
PSR
MT
PSR
MT
CTL
CTL
MO
MO
Heart
Heart
PSR
MT
PSR
MT
CTL
CTL
MO
MO
Supplemental Figure 2: Whole section fibrosis images for heart, kidney, and liver.
